# CCR5/CXCR3 antagonist TAK-779 prevents diffuse alveolar damage of the lung in murine model of the SARS-CoV-2-related acute respiratory distress syndrome

**DOI:** 10.1101/2023.11.14.567145

**Authors:** Aleksandr S. Chernov, Maksim V. Rodionov, Vitaly A. Kazakov, Karina A. Ivanova, Fedor A. Mesheryakov, Anna A. Kudriaeva, Alexander G. Gabibov, Georgii B. Telegin, Alexey A. Belogurov

## Abstract

The acute respiratory distress syndrome (ARDS) secondary to viral pneumonitis is one of the main causes of high mortality in patients with COVID-19 (novel coronavirus disease 2019) – ongoing SARS-CoV-2 infection, reached more than 0.7 billion registered cases. Recently we elaborated non-surgical and reproducible method of unilateral total diffuse alveolar damage (DAD) of the left lung in ICR mice – a publicly available imitation of the ARDS caused by SARS-CoV-2. Our data reads that two C-C chemokine receptor 5 (CCR5) ligands – macrophage inflammatory proteins (MIP) – (MIP-1α/CCL3) and (MIP-1β/CCL4) are upregulated in this DAD model up to three orders of magnitude compared to the background level. Here we showed that a nonpeptide compound TAK- 779, antagonist of CCR5/CXCR3, readily prevents DAD of the lung with a single injection of 2.5 mg/kg. Histological analysis revealed reduced peribronchial and perivascular mononuclear infiltration in the lung, and mononuclear infiltration of the wall and lumen of the alveoli in the TAK- 779-treated animals. Administration of the TAK-779 decreased 3-5-fold level of serum cytokines and chemokines in animals with DAD, including CCR5 ligands MIP-1α/β, MCP-1 and CCL5. Computed tomography revealed rapid recovery of the density and volume of the affected lung in TAK-779- treated animals. Our pre-clinical data suggest that TAK-779 is more effective than administration of dexamethasone or anti-IL6R therapeutic antibody tocilizumab, which brings novel therapeutic modality to TAK-779 and other CCR5 inhibitors recruited in ongoing clinical studies as a potential drugs for treatment of COVID19 and similar virus-induced inflammation syndromes.

**Abstract summary:** The pathogenesis of the SARS-CoV-2 infection is tightly linked with the cytokine storm resulting in the enormous release of cytokines and chemokines. Its clinical manifestation – the acute respiratory distress syndrome (ARDS), may be caused by self-sustaining hypersensitivity reactions leading to lung collapse even after virus clearance. Here we report that two macrophage inflammatory proteins, MIP-1α/CCL3 and MIP-1β/CCL4, seem to orchestrate mononuclear infiltration into the lungs during diffuse alveolar damage (DAD) in ICR mice – our murine model of ARDS caused by SARS-CoV-2. Inhibition of the C-C chemokine receptor 5 (CCR5) – parental receptor for MIP-1α and MIP-1β, by nonpeptide antagonist TAK-779 results in significant amelioration of DAD in terms of reduced mononuclear infiltration into the lung, suppressed cytokine storm and restored physiology of affected lung according to computed tomography data. We suggest that targeted inhibition of CCR5 should be further elucidated as safe and effective approach to overcome severe viral pneumonia in humans.

## Introduction

The socio-medical significance of the COVID-19 pandemic is hard to overestimate, as it has led to the death of more than 6.9 million people around the world. The treatment of severe COVID-19, which is accompanied by the development of acute respiratory distress syndrome or diffuse alveolar damage (ARDS/DAD) is still challenging [1]. Clinical signs of ARDS include a dysfunction of the alveolar epithelium and erupted gas exchange. An attempt to restore the condition to normal often leads to a fibroproliferative state [2]. The ARDS cause disability, morbidity and mortality, while current treatment is focused on basial patient support to give the lungs enough time to recover [3,4]. The course of ARDS is generally associated with a cytokine storm, consisting in the enormous release of cytokines and chemokines [5].

The immune response toward SARS-CoV-2 in humans occurs in several stages. Activation of type I interferon (IFN I) response induces the recruitment of macrophages, NK cells and neutrophils to the virus penetration site [6]. It is believed that resident alveolar macrophages are the first defensive line to fight the virus [7] and neutrophils may undergo TNF-dependent necroptosis among seriously ill patients [8,9]. During the adaptive immune response stage, CD4^+^ and CD8^+^ T cells spread and hyperactivate, which may lead to a lack of reactivity or cell death [10]. In addition, immune response modulation is actively driven by Th17 and Th22 subpopulations [5]. The level of IgG, IgM and IgA antibodies produced by B lymphocytes increases during COVID-19 infection [11]. B cells also release IL-6, which intensifies the cytokine storm [12]. As a result of cytokine release, the integrity of endotheliocytes is erupted and TxA2 is released, which in turn cause thrombosis [13]. The physiological effect of cytokines is maintained mainly through the activation of JAK/STAT signalling pathway. Level of cytokines, e.g. IL-6, positively correlates with the mortality of COVID-19 patients [14]. Fannning et al. showed that level of C-C/C-X-C chemokine ligands and C-reactive protein (CRP) are correlated with COVID-19 progression in patients treated by convalescent donor plasma [15]. As the cytokine storm escalates, apoptosis of the pulmonary epithelium increases, the blood–air barrier and vessels become damaged resulting in alveolar oedema and hypoxia [5]. High level of cytokines in the blood is also associated with bacterial superinfections in COVID-19 patients [8].

Current list of Food and Drug Administration (FDA)-approved COVID-19 therapeutics includes anti-interleukin-6 receptor monoclonal antibody Actemra (tocilizumab) [16,17], antiviral nucleotide analog Veklury (remdesivir) [18], inhibitor of Janus kinases JAK1 and JAK2 Olumiant (baricitinib) [19,20] and mixture of 3CL^pro^ protease inhibitor and inhibitor of HIV-1 protease CYP3A Paxlovid (nirmatrelvir and ritonavir) [21]. Therapy of coronavirus infection evolved in various directions. For example, remdesivir suppresses coronavirus replication by inhibiting RNA-dependent RNA polymerase [5]. Another approach to the COVID-19 treatment is administration of glucocorticosteroids. These immunosuppressors modify gene expression and trigger the synthesis of NF-kB inhibitors, reducing the production of IL-1 and IL-6 [5]. Treatment of COVID-19 patients was accomplished by monoclonal antibody olokizumab, which has high affinity to IL-6 and neutralizes this cytokine [22]. Furthermore, the anti-IL-17 monoclonal antibody netakimab, utilized for patients with psoriasis and ankylosing spondylitis, may be used for COVID-19 treatment [23]. Pharmacodynamically, netakimab effect is beneficial since IL-17 increases level of inflammatory mediators G-CSF, IL-6, IL-1β, TNFα, IL-8 and matrix metalloproteases [23]. Tocilizumab proved its clinical effectiveness in patients during the coronavirus pandemic [24]. One more approach of treatment is colchicine, an indirect IL-6 inhibitor, which is also applied in the treatment of coronary heart disease [25]. Colchicine suppresses the recruitment of neutrophils, inhibits cytoskeleton metabolism and opposes SARS-CoV-2 functionality in human cells [26]. Rodriguez et al. suggested IL-8 antagonists to be prospective agents to treat the severe coronavirus infection [27]. Indeed, convalescent plasma [28] and SARS-CoV-2 neutralizing antibodies [29,30] may be also regarded as direct agents for virus clearance.

Another pharmacological group, JAK inhibitors, such as tofacitinib, seems to be also perspective [31]. Baricitinib, a JAK1/JAK2 inhibitor, reduces the content of phosphorylated STAT proteins, which prevents the proinflammatory effect of IL-6, IL-12, IL-23 and IFN-γ [23]. It is also suggested that JAK-STAT pathway inhibition may ameliorate the condition during bacterial sepsis- induced ARDS [32]. The MAPK, RAS, and PI3K-AKT pathways may be potential targets in ARDS as well [32]. Metformin, used to treat type 2 diabetes, is also considered a promising drug that acts at different stages of the SARS-CoV-2 development (viral entry, viral replication etc.), but it has yet to be investigated [33]. Additionally, COVID-19 treatment can be carried out by HIFs pathway exposure [34]. Thus, preclinical trials of FG-4592 (Roxadustat) that suppressed PHDs were conducted. The drug activated HIFs and reduced viral burden and respiratory symptoms on the fourth day after infection [34]. Ewart et al. showed that small-molecule acylguanidine BIT225 prevents weight loss in SARS-CoV-2 infected K18 mice, suppresses virus reproduction and exhibits anti-inflammatory properties [35].

Summarizing, the global medical community is faced of creating and introducing into clinical practice new effective and safe methods of treating not only COVID-19, but similar virus-induced lung inflammation syndromes. Despite SARS-CoV-2 lost its pandemic status, the search for systemic and etiotropic treatment of ARDS should prepare humanity for possible future viral epidemics. The chemokine-driven migration of activated T cells and monocytes to the infection site is critical for the progression of COVID-19 [36,37]. Several reports showed that C-C/C-X-C chemokine ligands CCL3, CCL4 and CXCL-10 take part in the development of SARS-CoV-2 infection [5,38,39]. It is known that in the severe form of the disease, CXCR2 signalling is activated, while in the favourable course of the disease, a T helper “Th1-Th17” profile, marked by an upregulation of the CXCR3 pathway activator genes, is observed [27]. Recently, Dai et al. showed that H5N1 AIV-induced inflammatory lung injury is driven by infiltrating inflammatory macrophages with massive viral replication and emerging interaction of cell populations through various chemokines, including CCL4 [40]. Thus, possible way to treat coronavirus infection is to target chemokine receptors. Among the three anti-chemokine drugs (leronlimab, maraviroc and cenicriviroc), the latter seems to be the most optimal, since it, by inhibiting CCR2 and CCR5 pathways, precludes the pulmonary and vascular sequelae associated with COVID-19 [41].

The TAK-779 is also a potent and selective nonpeptide antagonist of CCR5/CXCR3 [42], with a K_i_ of 1.1 nM. The CCR5-related cognate ligands include CCL3, CCL4 (also known as MIP 1α and 1β, respectively), and CCL3L1. The CCR5 also interacts with CCL5 (a chemotactic cytokine protein, also known as RANTES). TAK-779 effectively and selectively inhibits R5 HIV-1 in MAGI-CCR5 cells with EC_50_ of 1.2 nM [43]. In dosage of 10 mg/kg per day TAK-779 significantly prolongs the allograft survival of the rat intestinal transplantation model. It inhibits migration of T cells but not its proliferation. TAK-779 also decreases the number of CD4^+^ as well as CD8^+^ T cells in spleen, blood and recipient mesenteric lymph nodes (MLN) [44]. Zhu et al. demonstrate that TAK-779 in dosage of 150 µg per mouse suppresses the development of experimental autoimmune encephalomyelitis (EAE) in myelin oligodendrocyte glycoprotein (MOG)-immunized C57BL/6 mice. It decreases the infiltration of CXCR3- and CCR5-bearing leukocytes into the spinal cord [45]. TAK-779 ameliorates pulmonary granulomatosis in C57BL/6 mice and diminishes the pool of CXCR3^+^CD4^+^ and CCR5^+^CD4^+^ T lymphocytes in the bronchoalveolar lavage fluid [46]. Tokuyama et al. showed that TAK-779 decreases the recruitment of monocytes/macrophages, thereby precluding dextran sodium sulfate-induced colitis in C57BL/6 mice [47]. TAK-779 also impedes the progression of chronic vasculopathy, fibrosis, and cellular infiltration by reducing the number of CD4, CD8, and CD11c- positive cells recruited to the transplanted allografts [48]. Furthermore, it was shown that in a Pan02 murine tumor model, a TAK-779 induced disruption of CCR5/CCL5 pathway led to a decreased Treg migration to the tumor [49]. Likewise, TAK-779 inhibits the homing of microglia in response to scrapie infection *in vitro* and *in vivo* [50].

The main restriction of the COVID-19 modeling in mice is that the SARS-CoV-2 does not bind to the mouse ACE2 (mACE2) [51]. Thus, various artificial models are used to study the effects of coronavirus infection. Yinda et al. reported that the administration of 10^4^ TCID_50_ or 10^5^ TCID_50_ SARS-CoV-2 led to 80% and absolute lethality, respectively, in K18-hACE2 mice, as well as to a mild form of the disease in mice who received 10^2^ TCID_50_ SARS-CoV-2 [52]. Nevertheless, the disadvantage of this model is the possible fatal outcome after a fulminant SARS-CoV-2 brain infection [53,54]. Two additional major obstacles of animal models, involving SARS-CoV-2 inoculation, are absence of prolonged monitoring due to the high mortality rate and generally high variability between individual animals in terms of cytokine levels and other immunological indicators. These features limit statistical analysis and yield binary “yes or no” result of the experimental therapy. Here we assessed therapeutic effect of dexamethasone, tocilizumab and TAK- 779 on the course of the ARDS in non-lethal, highly reproducible ICR DAD murine model mimicking SARS-CoV-2 infection [55] using histological evaluation, cytokine profiling and computed tomography analysis.

## Results

### Treatment by TAK-779 significantly reduces mononuclear infiltration in the lung in murine model of the SARS-CoV-2-related acute respiratory distress syndrome

The DAD was induced in ICR mice by a single instillation into the left lung of the mixture consisting of LPS from *Salmonella enterica* and α-galactosylceramide. On the 7th day of monitoring, a similar pathomorphological finding was observed in all test groups: total or subtotal atelectasis of the left lobe with pronounced neutrophilic and mononuclear infiltration into the walls and lumen of the alveoli, moderate focal peribronchial and perivascular infiltration (**Figure 1A**). The right lobes were intact in all cases.

**Figure 1.**
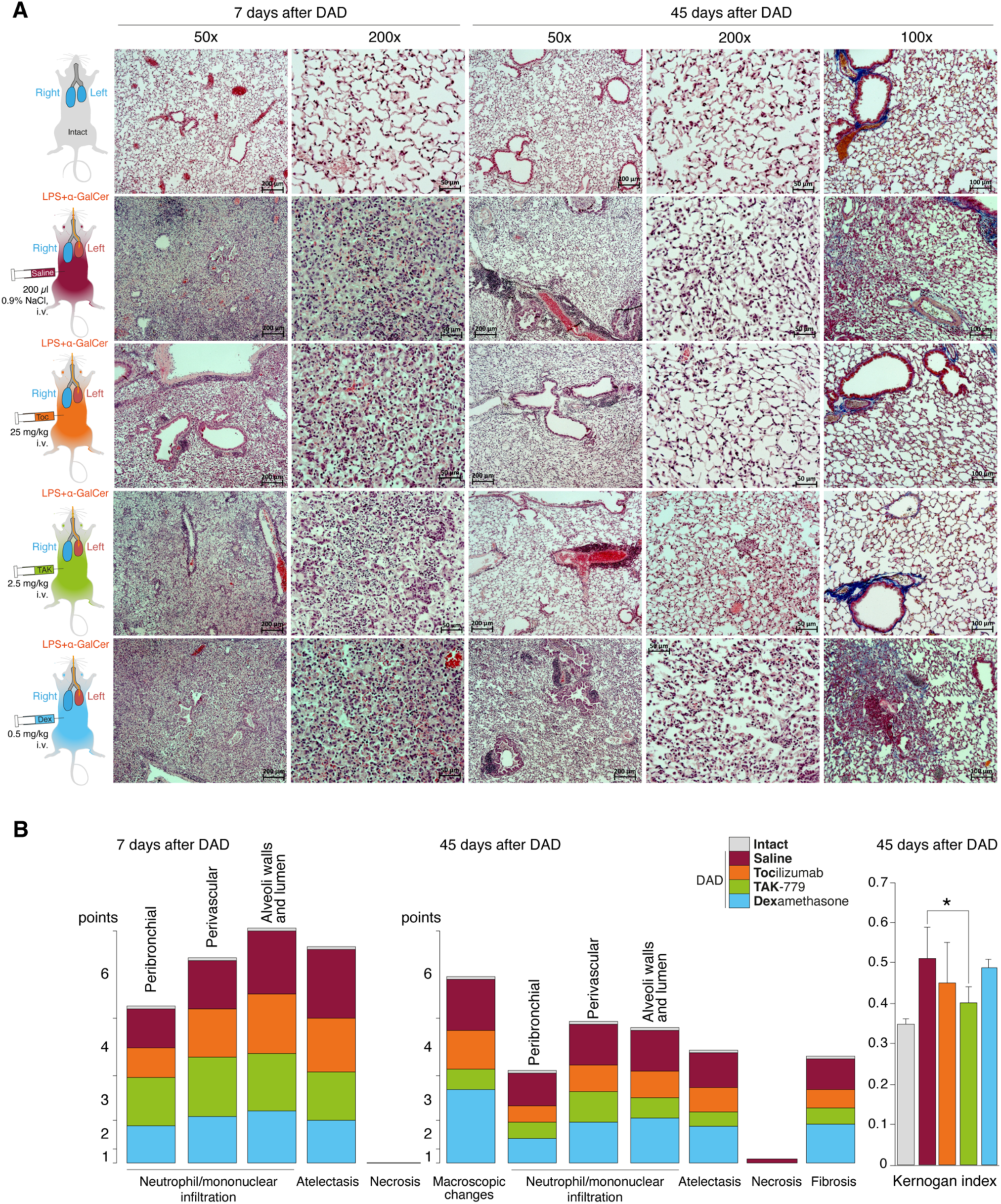
Reduced peribronchial and perivascular mononuclear infiltration in the lung, and mononuclear infiltration of the wall and lumen of the alveoli in the TAK-779-treated ICR mice with DAD. (**A**) Histological analysis of the left lobe of the lungs of ICR mice with DAD on day 7^th^ and 45^th^ in comparison with unexposed animals (grey) and mice treated by saline (control, red), tocilizumab (orange), TAK-779 (green) and dexamethasone (blue). Stained with hematoxylin and eosin and by Mallory method (right column). Magnification 50x, 100x and 200x. (**B**) Semi-quantitative score (0-5 points) of peribronchial and perivascular mononuclear infiltration, mononuclear infiltration into the wall and lumen of the alveoli, atelectasis, necrosis and fibrosis on 7^th^ and 45^th^ day after DAD induction in unexposed (intact, grey), untreated (saline, red) and mice with DAD treated by tocilizumab (orange), TAK-779 (green) and dexamethasone (blue). Kernogan index after 45 days post DAD induction in test groups is shown on right. Bars represent standard deviation. Statistically significant difference with non-treated animals with DAD (saline, red) is marked by asterisk.

The degree of atelectasis of the left lobe of the lungs was minimal in the group of animals with DAD receiving dexamethasone and maximal in case of saline administration (placebo) (**Figure 1B**, **Table 1**). In animals treated with dexamethasone the degree of neutrophil/mononuclear infiltration of the walls and lumen of the alveoli was estimated at 3.6 points, focal perivascular neutrophil/mononuclear infiltration corresponded to 3.3 points (**Figure 1B**, **Table 1**). The degree of focal peribronchial neutrophil and mononuclear infiltration was minimal among animals receiving tocilizumab and maximal among animals receiving TAK-779 (**Figure 1B**, **Table 1**). Focal perivascular neutrophil/mononuclear infiltration was also the highest in TAK-779-treated group (**Figure 1B**, **Table 1**). Statistically significant difference between treated and non-treated animals was observed in the rate of the atelectasis in groups of mice received TAK-779 and tocilizumab.

**Table 1.**
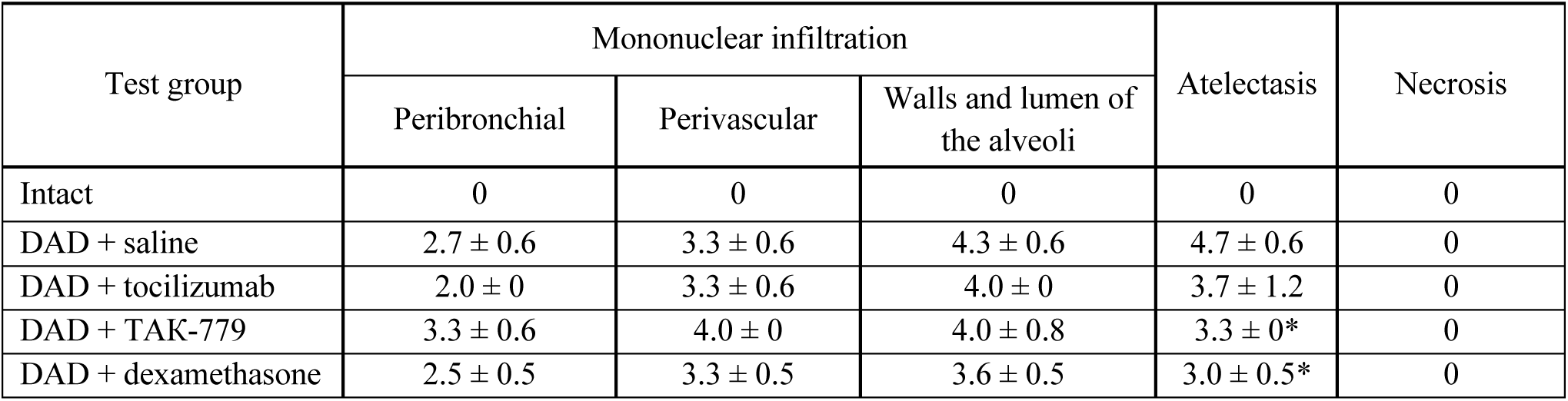
Left lung damage score on 7^th^ day after DAD induction in ICR mice treated with saline, tocilizumab, ТАК-779 and dexamethasone. Statistically significant difference with non-treated animals (saline) is marked by asterisk.

| Test group | Mononuclear infiltration |  |  | Atelectasis | Necrosis |
| --- | --- | --- | --- | --- | --- |
|  | Peribronchial | Perivascular | Walls and lumen of the alveoli |  |  |
| Intact | 0 | 0 | 0 | 0 | 0 |
| DAD + saline | 2.7 ± 0.6 | 3.3 ± 0.6 | 4.3 ± 0.6 | 4.7 ± 0.6 | 0 |
| DAD + tocilizumab | 2.0 ± 0 | 3.3 ± 0.6 | 4.0 ± 0 | 3.7 ± 1.2 | 0 |
| DAD + TAK-779 | 3.3 ± 0.6 | 4.0 ± 0 | 4.0 ± 0.8 | 3.3 ± 0* | 0 |
| DAD + dexamethasone | 2.5 ± 0.5 | 3.3 ± 0.5 | 3.6 ± 0.5 | 3.0 ± 0.5* | 0 |

Total lesion of the left lobe with a significant reduction in its volume and total hypoventilation in all tested groups was observed 45 days after DAD induction (**Figure 1A**). Moderate focal perivascular and insignificant focal peribronchial mononuclear infiltration was detected. The walls and lumen of the alveoli were also diffusely infiltrated by mononuclear cells (**Figure 1B**, **Table 2**). The lumen of a significant part of the bronchi was partially or completely obstructed. Mononuclear infiltration of the walls and lumen of the alveoli, as well as signs of hypoventilation in case of dexamethasone administration showed negative dynamics in comparison with non-treated group (**Figure 1B**, **Table 2**). The latter was confirmed by the average estimated score of macroscopic changes (5.0) in the left lobe of the animals receiving dexamethasone (**Figure 1B**, **Table 2**). The fibrous changes in the left lobe also had a negative trend and were estimated at 2.7 points. The Kernogan index of dexamethasone-treated mice was comparable with the group of animals receiving saline (0.49 ± 0.02) (**Figure 1B**).

**Table 2.** Left lung damage score on 45^th^ day after DAD induction in ICR mice treated with saline, tocilizumab, ТАК-779 and dexamethasone. Statistically significant difference with non-treated animals (saline) is marked by asterisk.

| Test group | Mononuclear infiltration |  |  | Macroscopic changes | Atelectasis | Necrosis | Fibrosis |
| --- | --- | --- | --- | --- | --- | --- | --- |
|  | Peribronchial | Perivascular | Walls and lumen of the alveoli |  |  |  |  |
| Intact | 0 | 0 | 0 | 0 | 0 | 0 | 0 |
| DAD + saline | 2.1 ± 0.7 | 2.9 ± 0.6 | 2.9 ± 0.5 | 3.5 ± 0.6 | 2.4 ± 0.8 | 0.29 ± 0.08 | 2.0 ± 0 |
| DAD + tocilizumab | 1.2 ± 0.5* | 1.8 ± 0.5* | 1.8 ± 0.8* | 2.6 ± 0.4 | 1.6 ± 0.3 | 0 | 1.3 ± 0.3* |
| DAD + TAK-779 | 1.0 ± 0.3* | 2.0 ± 0* | 1.3 ± 0.4* | 1.3 ± 0.4* | 1.0 ± 0* | 0 | 1.0 ± 0* |
| DAD + dexamethasone | 1.7 ± 0.8 | 2.8 ± 0.7 | 3.0 ± 0.8 | 5.0 ± 0* | 2.5 ± 0.1 | 0 | 2.7 ± 0.5 |

Administration of tocilizumab resulted in significant improvement in the pathomorphological finding on the 45^th^ day of observation in the left lobe of the lungs for all key indicators (**Figure 1B**, **Table 2**). The average score of mononuclear infiltration of the walls and lumen of the alveoli was estimated at 1.8 points, focal peribronchial and perivascular mononuclear infiltration – at 1.2 and 1.8 points, respectively. In addition, the left lobe of the lungs participated in gas exchange as areas of atelectasis and fibrous changes were estimated on average at 1.6 (macroscopic score – 2.6) and 1.3 points, respectively (**Table 2**). The Kernogan index had more favorable values (0.45 ± 0.10) compared to groups of animals treated with saline and dexamethasone (**Figure 1B**).

The most pronounced improvement on the 45^th^ day of observation was detected in tested group treated by TAK-779 (**Figure 1B**, **Table 2**). Macroscopic changes in the left lobe of the lungs corresponded to 1.3. Atelectasis according to histological data was estimated as 1.0. The degree of mononuclear infiltration of the walls and lumen of the alveoli was determined at 1.3 points, focal peribronchial and perivascular mononuclear infiltration – at 1.0 and 2.0 points, respectively. The degree of fibrous changes in the left lobe was 1.0 points. The Kernogan index was also the most favorable in comparison with all the analyzed groups – 0.40 ± 0.04 (**Figure 1B)**.

### Treatment by TAK-779 suppresses development of cytokine storm in murine model of the SARS-CoV-2-related acute respiratory distress syndrome

We next measured cytokine profiles in treated and non-treated ICR mice with induced DAD and compared it with unexposed animals (**Figure 2A**). Our data suggest that during first 3 hours almost all cytokines and chemokines were significantly elevated in the plasma of mice with induced DAD compared to intact animals (**Figure 2B**). Level of the IL-5, IL-12p40, tumor necrosis factor (TNF) and interferon-gamma (IFNγ) was increased 2-5 fold, level of IL-1a, IL-6, IL-9, IL-10, IL-12p70 and regulated on activation, normal T cell expressed and secreted (RANTES/CCL5, chemokine C-C motif ligand 5) was upregulated up to 10 times, level of IL-4, monocyte chemoattractant protein 1 (MCP- 1/CCL2, chemokine C-C motif ligand 2), keratinocyte chemoattractant (KC) and granulocyte colony- stimulating factor (G-CSF) was elevated up to two orders of magnitude in comparison with unexposed animals. Two CCR5 ligands macrophage inflammatory proteins (MIP) – (MIP-1α/CCL3) and (MIP-1β/CCL4) were upregulated up to three orders of magnitude compared to background level. Administration of dexamethasone did not alter cytokine and chemokine profile mice with DAD except level of TNF, which decreased twice in comparison with untreated animals. Injection of tocilizumab and TAK-779 resulted in similar, 3-5-fold downregulation of majority of cytokines except IL-6, MCP-1 and G-CSF (**Figure 2B**).

**Figure 2.**
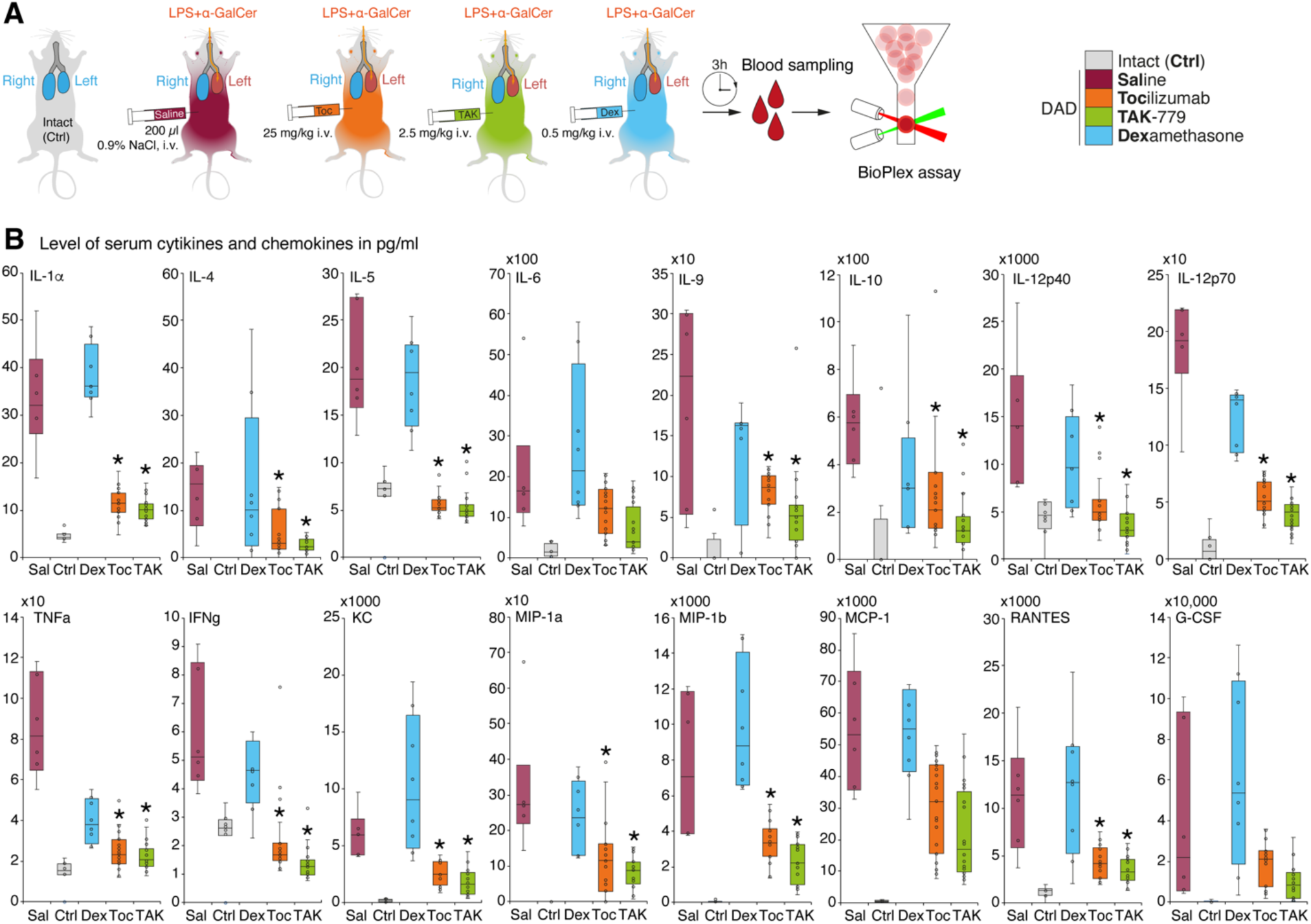
Administration of the TAK-779 decreases 3-5-fold level of serum cytokines and chemokines in ICR mice with DAD. **(A)** Study design. Blood was collected after 3 hours post DAD induction and level of cytokines and chemokines was estimated by multiplex immunoassay. **(B)** Level of plasma cytokines and chemokines (pg/ml) in ICR mice 3 hours after DAD induction treated by tocilizumab (orange), TAK-779 (green) and dexamethasone (blue) in comparison with unexposed (grey) and non-treated animals with DAD (red). Bars represent median, interquartile range with standard deviation. Granulocyte colony-stimulating factor (G-CSF), keratinocyte chemoattractant (KC), tumor necrosis factor (TNF), interferon gamma (IFNγ), regulated on activation, normal T cell expressed and secreted (RANTES/CCL5), monocyte chemoattractant protein 1 (MCP-1/CCL2), and macrophage inflammatory proteins (MIP) – MIP-1α and MIP-1β. Statistically significant difference with non-treated animals with DAD (saline, red) is marked by asterisk.

### Treatment by TAK-779 prevents collapse of the injured lung in murine model of the SARS-CoV-2-related acute respiratory distress syndrome

A dynamic assessment of the volume of pulmonary lobes and its average density in Hounsfield units of the ICR mice from test groups was performed using a CT scanner MRS*CT/PET on 7^th^, 14^th^, 30^th^ and 45^th^ day after DAD induction (**Figure 3A**). Generally, we observed development of the specific pattern in the left lung in all groups: total or subtotal consolidation of the lung volume, less often mosaic-located areas of ground-glass opacities, alveolar consolidation, and rare areas of the intact lung tissue (**Figure 3B**).

**Figure 3.**
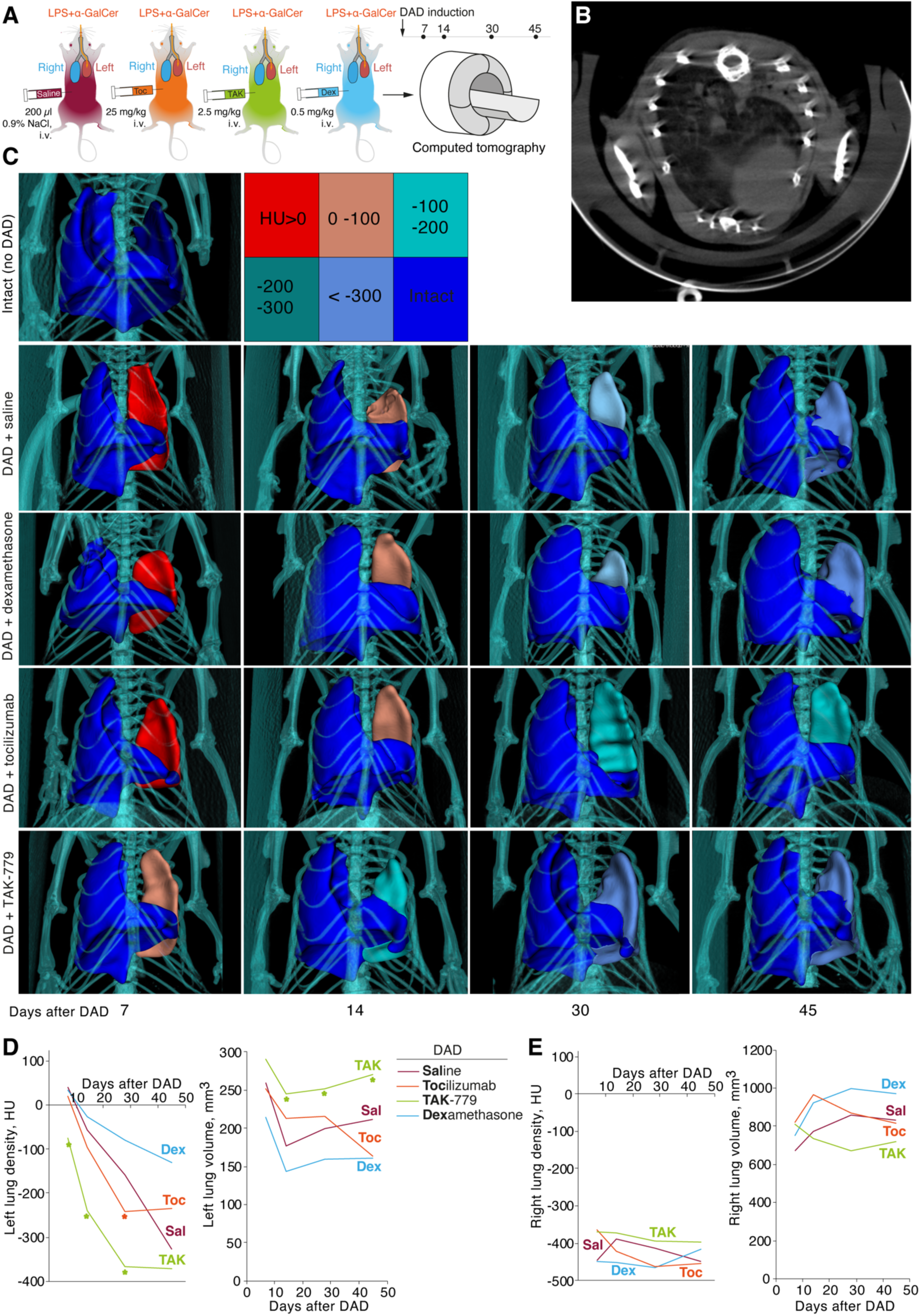
Treatment by TAK-779 induces fast recovery of injured lung in ICR mice with DAD. (**A**) Study design. Computed tomography (CT) was performed after 7, 14, 30 and 45 days post DAD induction. (**B**) Representative CT visualization of mice lungs after DAD induction. Subtotal consolidation of the lung volume, less often mosaic-located areas of ground-glass opacities, alveolar consolidation, and rare areas of the intact lung tissue were observed. (**C**) Representative 3D CT reconstruction of the murine lungs from different groups. Color legend for Hounsfield units is shown. Density (HU) and volume (mm^3^) of murine left (**D**) and right (**E**) lung of non-treated (Sal, red) and tocilizumab (Toc, orange), TAK-779 (TAK, green) and dexamethasone (Dex, blue)-treated animals. Statistically significant difference with non-treated animals with DAD is marked by asterisk.

Seven days after DAD induction, density of the affected left lung of the non-treated mice and mice treated by tocilizumab and dexamethasone was 40.0 HU, 20.4 HU and 33.4 HU, respectively (**Figure 3С,D**). In mice treated by TAK-779 the density of the left lung was significantly lower and equaled to -75 HU (**Figure 3C,D**). Volume of the affected lung 14 days after DAD induction decreased less than 40% in the group of mice treated by TAK-779, whereas non-treated animals and dexamethasone-treated mice showed 60-70% collapse of the left lung volume. Further monitoring of the density and volume of the affected lung in test groups revealed rapid recovery of the injured lung in TAK-779- treated mice, minimal beneficial effect of the tocilizumab and negative influence of the dexamethasone administration. Treatment by TAK-779 firstly restored the caudal segments of the lung and further normalized the segments of the apex. Parameters of the non-affected right lung of mice from all test groups (**Figure 3E**) were equal to those of the intact mice.

## Discussion

In majority of clinical cases innate and adaptive immunity overcomes SARS-CoV-2 infection, however, some patients develop a severe late stage, which promotes the cytokine storm and accelerates the deterioration of patients to the ARDS/DAD [56]. The main pathological effect may be caused by hypersensitivity reactions rather than virus itself, as study by Zhenfei et al. reported that formaldehyde-inactivated SARS-CoV-2 induces ARDS in human ACE2-transgenic mice [57].

Dexamethasone has evident anti-inflammatory effect and is widely used as an auxiliary treatment for viral pneumonia [58]. Low and moderate doses of dexamethasone decrease the mortality rate in patients with a severe form of the COVID-19. However, it is not recommended for patients with mild symptoms [59]. In patients hospitalized with COVID-19, the use of dexamethasone resulted in lower 28-day mortality among those who were receiving either invasive mechanical ventilation or oxygen alone but not among those receiving no respiratory support [60]. Interestingly, it was demonstrated that 6 mg dexamethasone (7-10 days) was less effective in accelerating recovery and reducing severity markers (CRP, D-dimer and LDH) than high-dose methylprednisolone (3 days) followed by prednisone (2 weeks) [61]. Moreover, in other clinical study 60-day mortality was not reduced when high-dose dexamethasone was prescribed in patients with COVID-19-related acute hypoxemic respiratory failure [62]. In another randomized controlled trial, it was found that 28-day mortality was increased in hospitalized patients with COVID-19 pneumonia receiving high doses of dexamethasone (20 mg q.d.), in contrast to patients receiving 6 mg of dexamethasone q.d. [63].

On the COVID-19 model in hamsters, dexamethasone prevented inflammation and helped preserve the integrity of the lungs [64]. Moreover, in hamsters dexamethasone did not accelerate the virus replication and diminished the number of inflammatory mediators [64]. Clinically, the latter effect is important for preventing pulmonary edema and enhancing gas exchange [64]. In SARS- CoV-2 infected rhesus macaques, a dose of 0.01 mg/kg inhaled nanoDEX (engineered neutrophil nanovesicles to deliver dexamethasone) reduced lung inflammation and preserved their integrity better compared to intravenous administration of 0.1 mg/kg dexamethasone [65]. However, in other hamster COVID-19 model, dexamethasone administration contributed to a decrease in serum- neutralizing antibody and RBD-specific antibody titer titers, which led to a minor growth of viral replication [66].

Our data suggest that dexamethasone at least in our DAD model and dosage regime did not have any resulted beneficial effect on ARDS pathology. It showed some non-statistically significant effect revealed by histological analysis in the early period of ARDS but failed to show any efficacy according to other evaluation techniques. One may suggest that murine models may be inappropriate for testing of dexamethasone therapeutic potential as Xu et al. showed that dexamethasone treatment in dosage of 2.5 mg/kg from days 3-14 post inoculation has no beneficial effect on ARDS in mice caused by the H5N1 virus [67].

Tocilizumab was shown to be effective treatment in patients with severe COVID-19 [68]. It has a positive effect on improving immune damaging, lung functional injuries and arterial oxygen saturation. In addition, it was shown that tocilizumab reduces the number of HDL-1 subfractions of cholesterol, phospholipids, and Apo A2, and increases the level of LDL-5, HDL-4, IDL, VLDL-1, and VLDL-2, which helps to partially restore the indicators changed due to coronavirus infection [69]. However, in randomized trial involving hospitalized patients with severe COVID-19 pneumonia, the use of tocilizumab did not result in significantly better clinical status or lower mortality than placebo at 28 days [70]. Moreover, in other randomized clinical trial, it was shown that after the tocilizumab administration, the risk of intubation or death, disease condition, or time to discontinuation of supplemental oxygen did not significantly change in patients with COVID-19 [71]. In another clinical trial, it was found that 400 mg tocilizumab did not improve hypoxemia on day 14 or 28 and did not change ventilator-free survival on day 14 among patients with COVID-19 [72]. Finally, the following clinical trial revealed that tocilizumab (8 mg/kg) may elevate mortality and its administration did not lead to better clinical outcomes at 15 days [73]. In a murine model of COVID- 19, it was found that the use of interleukin-6 receptor blockers does not change the content of IL-1 and TNF, although it diminishes neutrophil infiltration [74].

We showed that tocilizumab significantly decreased release of cytokines and chemokines and inhibited mononuclear infiltration during DAD development. Despite several reports claiming effect of tocilizumab in various experimental murine models [75–78], study by Lokau et al. [79] showed that tocilizumab does not block IL-6 signaling in murine cells. These data may explain absence of tocilizumab effect on complex injured lung restoration in our study visualized by computed tomography.

COVID-19 also may be treated with a leronlimab – humanized monoclonal antibody to CCR5 (PRO 140), which binds to the extracellular loop 2 domain and the N-terminus of CCR5 [80]. Interestingly, healthy *rhesus macaques* receiving 10 mg/kg or 50 mg/kg leronlimab injection had elevated CCR5^+^CD4^+^ T cell counts [81]. Furthermore, this drug was administered subcutaneously to COVID-19 patients on the 0 and 7 days of the study, which led to a decrease in the inflammatory mediator IL-6 concentration, recovery of the CD4/CD8 ratio and a reduction in plasma viremia (pVL) [80]. Another case study also showed a decrease in the first and a recovery of the second indicator [82]. In “long COVID-19”, leronlimab administrated 700 mg s.c. weekly also normalized the immune downmodulation [83].

Files et al. reported that CCR2 and CCR5 blocker cenicriviroc reduces the infiltration of monocytes and lymphocytes into the lung tissue [41] and prevents the replication of SARS-CoV-2 in VeroE6/TMPRSS2 cells by inhibiting virus-dependent cell destruction (EC_50_ = 19 µM) [84]. This effect may not have a direct impact on the virus, but the possible restriction of myeloid-derived suppressor cells may increase the pool of effector lymphoid B and T cells that will exhibit antiviral exposure in an indirect way [41]. In one clinical trial, сenicriviroc (100 and 200 mg) was effective and well-tolerated by HIV-1-infected patients [85]. Thus, a growth in the concentration of cenicriviroc contributed to the improvement of virological outcomes [85]. In another clinical trial, cenicriviroc (150 mg) ameliorated fibrosis, prevented the progression of steatohepatitis and diminished the indicators of systemic inflammation in patients with non-alcoholic steatohepatitis [86]. In the third clinical trial in patients with the same disease, cenicriviroc (150 mg) also precluded systemic inflammation and fibrosis by reducing C-reactive protein, IL-6, IL-1β, and fibrinogen levels [87]. However, a randomized clinical trial revealed that hospitalized COVID-19 patients treated with cenicriviroc did not recover faster than those patients who received a placebo [88]. Another low- weight CCR5 antagonist, maraviroc, was also applied for COVID-19 treatment. This drug is quite promising due to its low protein binding and high bioavailability [89]. At the same time, maraviroc strongly inhibits SARS-CoV-2 Mpro and supresses the coronavirus infection development [89]. In “long COVID-19”, the combination of maraviroc and pravastatin improved clinical indicators among 18 patients in a case study [90]. As of today, two clinical trials (NCT04435522 and NCT04710199) have been completed and two (NCT04441385 and NCT04475991) have been terminated.

Nevertheless, in African green monkey kidney cell model of COVID-19, it was demonstrated that the administration of maraviroc reduces the viral load by precluding membrane fusion, which affects the reproduction and dissemination of the coronavirus [91]. In the same study, it was revealed that maraviroc supresses S-protein transport to the extracellular surface of the cell [91].

Our data uniquely reads that selective CCR5/CXCR3 inhibitor TAK-779 is highly efficient in the prevention of the ARDS in our murine DAD model. Its therapeutic effect is reasoned by a milder course of the inflammatory process in the lungs with early and effective involvement of the affected lung lobe in systemic gas exchange. These suggestions are confirmed by low degree of fibrosis and volume loss of the injured lung, decreased mononuclear infiltration and rate of density restoration among animals treated with TAK-779. More favorable Kernogan index observed in this group is as an important quantitative indicator of the capacity of small circle vessels and, as a consequence, an indicator of the load on systemic blood flow. Our study indicates that TAK-779 can effectively alleviate DAD and lung collapse in mice by its direct inhibitory effects on inflammation, cytokine release and immune cell migration. We suggest that several ongoing clinical studies of CCR5 inhibitors may provide not only promising agents for cancer treatment but also novel therapeutic modalities for amelioration of COVID-19 and similar virus-induced inflammation syndromes.

## Materials and methods

### Ethics statement

ICR male mice SPF-category with an average weight of 37.4 ± 1.4 g were used. All animals were housed under standard conditions in the Animal breeding facility of BIBCh RAS (the Unique Research Unit Bio-Model of the IBCh RAS; the Bioresource Collection – Collection of SPF- Laboratory Rodents for Fundamental, Biomedical and Pharmacological Studies, Сontract 075-15- 2021-1067). All experiments and manipulations with animals were approved by the institutional animal care and use committee (IACUC № 831/22 from 12/04/22).

### Reagents

LPS (*Salmonella enterica*) was purchased from Millipore Sigma (USA), α-galactosylceramide was from Avanti Polar Lipids (USA), propofol was from Hana Pharmaceutical, Co. Ltd. (Republic of Korea), TAK-779 was from MedChemExpress (USA), tocilizumab was from Roshe (Japan).

### DAD modeling in ICR mice

DAD was induced by a single instillation into the left lung of the mixture consisting of 100 μl (1 mg/ml) LPS from *Salmonella enterica* and 100 μl (50 μg/ml) α-galactosylceramide with an intravenous catheter 20G. As premedication intravenous bolus into the lateral tail vein at a dose of 20 mg/kg a short-acting hypnotic agent propofol was used.

### Administration and doses of test substances

The animals were randomly assigned to five groups with 20 animals in each group: (1) – control, intact animals; (2) – induction of DAD and intravenous injection of physiological saline (200 µl 0.9% NaCl) as a placebo; (3) – induction of DAD and intravenous injection of tocilizumab (25 mg/kg); (4) – induction of DAD and intravenous injection of TAK-779 (2.5 mg/kg); (5) – induction of DAD and intravenous injection of dexamethasone (0.5 mg/kg). Animals were monitored daily for 45 days.

### Computed tomography of the lungs

CT scanner MRS*CT/PET (MR Solution, UK) was used with following parameters: energy 40 kVp, exposure 100 ms, current 1 mA, stepping angle 1°. During the imaging, the animals were anesthetized with a 2% mixture of isoflurane and air, at a temperature of +37°С. The CT images were processed using the VivoQuant software (Invicro, UK). The lung volume and the average density of left (exposed) and right (control) lungs were measured in the automatic mode (using -400 to 100 Hounsfield units as the cut-off density). The volume was expressed in mm^3^ and the density in Hounsfield units (HU).

### Plasma chemokines and cytokines measurement

Mouse blood for EDTA plasma was collected after 3 hours post DAD induction. Bio-Plex Pro Mouse Cytokine Panel 33-Plex (Bio-Rad, USA) was used to measure the levels of chemokines and cytokines. Plasma samples diluted at 1:3 (50 μl) were incubated with magnetic beads, washed up, and then incubated with detecting antibodies and SA-PE according manufacturer’s instructions. Data were obtained using Luminex 200 analyzer and analyzed using the xPONENT software.

### Histological examination

Five animals from each group were sacrificed at day 7 and 45 and histological studies were performed, as described previously [55,92]. Briefly, the lungs were filled with 10% solution of neutral formalin, embedded in paraffin and 4–5 μm width sections were stained with hematoxylin and eosin. The degree of fibrosis was examined on histological preparations stained by the Mallory method. Histological analysis of the lungs assessed the following morphological signs: peribronchial and perivascular mononuclear infiltration, infiltration of the walls and lumen of the alveoli by mononuclears, atelectasis, the presence or absence of necrosis foci and level of fibrosis. The severity of various inflammatory phenomena in the lungs and the degree of pneumofibrosis were evaluated by a semi–quantitative method (in points) according to the 5-score scale [92]. The Kernogan index in the blood vessels of the left lobe of the lungs was evaluated as the ratio of the thickness of the vascular wall to the radius of the vessel lumen as an important indicator of the throughput of the microcirculatory bed of the small circulatory circle using ZEN 2.6 lite software (Carl Zeiss, Germany).

### Statistical analysis

Data are presented as mean ± standard deviation. Differences between treatment and control were tested for significance with SigmaPlot software (Systat Software Inc, Berkshire, UK), using Student’s t-test.

## Acknowledgements

This work was carried out within the framework of the Russian Ministry of Education and Science project No. 075-15-2021-1049.

## Author contributions

Conceptualization, A.S.C., G.B.T., A.G.G. and A.A.B.; methodology, M.V.R., V.A.K., A.A.K.; software, ; validation, formal analysis, A.A.B.; investigation, A.A.K., M.V.R., V.A.K.; resources,; data curation, A.A.K., F.A.M.; writing – original draft preparation, A.S.C., K.A.I., A.A.B.; writing – review and editing, A.S.C., K.A.I., A.A.B.; visualization, A.A.B.; supervision, A.S.C., G.B.T., A.G.G. and A.A.B.; project administration, A.S.C., G.B.T.; funding acquisition, A.S.C., G.B.T., A.G.G. All authors have read and agreed to the published version of the manuscript.

## Notes

### Competing Interest Statement

The authors have declared no competing interest.

## References

1. Ntatsoulis K, Karampitsakos T, Tsitoura E, Stylianaki E-A, Matralis AN, Tzouvelekis A, et al. Commonalities Between ARDS, Pulmonary Fibrosis and COVID-19: The Potential of Autotaxin as a Therapeutic Target. Front Immunol. 2021;12: 687397. doi:10.3389/fimmu.2021.687397

2. Fears AC, Beddingfield BJ, Chirichella NR, Slisarenko N, Killeen SZ, Redmann RK, et al. Exposure modality influences viral kinetics but not respiratory outcome of COVID-19 in multiple nonhuman primate species. PLoS Pathog. 2022;18: e1010618. doi:10.1371/journal.ppat.1010618

3. Wick KD, McAuley DF, Levitt JE, Beitler JR, Annane D, Riviello ED, et al. Promises and challenges of personalized medicine to guide ARDS therapy. Crit Care. 2021;25: 404. doi:10.1186/s13054-021-03822-z

4. Luo X, Lv M, Wang X, Long X, Ren M, Zhang X, et al. Supportive care for patient with respiratory diseases: an umbrella review. Ann Transl Med. 2020;8: 621. doi:10.21037/atm-20-3298

5. Martonik D, Parfieniuk-Kowerda A, Starosz A, Grubczak K, Moniuszko M, Flisiak R. Effect of antiviral and immunomodulatory treatment on a cytokine profile in patients with COVID-19. Front Immunol. 2023;14: 1222170. doi:10.3389/fimmu.2023.1222170

6. Tamir H, Melamed S, Erez N, Politi B, Yahalom-Ronen Y, Achdout H, et al. Induction of Innate Immune Response by TLR3 Agonist Protects Mice against SARS-CoV-2 Infection. Viruses. 2022;14. doi:10.3390/v14020189

7. Kumar R, Aktay-Cetin Ö, Craddock V, Morales-Cano D, Kosanovic D, Cogolludo A, et al. Potential long-term effects of SARS-CoV-2 infection on the pulmonary vasculature: Multilayered cross-talks in the setting of coinfections and comorbidities. PLoS Pathog. 2023;19: e1011063. doi:10.1371/journal.ppat.1011063

8. Mairpady Shambat S, Gómez-Mejia A, Schweizer TA, Huemer M, Chang C-C, Acevedo C, et al. Hyperinflammatory environment drives dysfunctional myeloid cell effector response to bacterial challenge in COVID-19. PLoS Pathog. 2022;18: e1010176. doi:10.1371/journal.ppat.1010176

9. Schweizer TA, Mairpady Shambat S, Vulin C, Hoeller S, Acevedo C, Huemer M, et al. Blunted sFasL signalling exacerbates TNF-driven neutrophil necroptosis in critically ill COVID-19 patients. Clin Transl Immunol. 2021;10: e1357. doi:10.1002/cti2.1357

10. Alberca RW. Immune Response to COVID-19. In: Baddour MM, editor. Rijeka: IntechOpen; 2021. p. Ch. 6. doi:10.5772/intechopen.98964

11. Ruhl L, Pink I, Kühne JF, Beushausen K, Keil J, Christoph S, et al. Endothelial dysfunction contributes to severe COVID-19 in combination with dysregulated lymphocyte responses and cytokine networks. Signal Transduct Target Ther. 2021;6: 418. doi:10.1038/s41392-021-00819-6

12. Upasani V, Rodenhuis-Zybert I, Cantaert T. Antibody-independent functions of B cells during viral infections. PLoS Pathog. 2021;17: e1009708. doi:10.1371/journal.ppat.1009708

13. Conti P, Pregliasco FE, Calvisi V, Caraffa A, Gallenga CE, Kritas SK, et al. Monoclonal antibody therapy in COVID-19. Journal of biological regulators and homeostatic agents. Italy; 2021. pp. 423–427. doi:10.23812/Conti_Edit_35_2_1

14. Stebbing J, Sánchez Nievas G, Falcone M, Youhanna S, Richardson P, Ottaviani S, et al. JAK inhibition reduces SARS-CoV-2 liver infectivity and modulates inflammatory responses to reduce morbidity and mortality. Sci Adv. 2021;7. doi:10.1126/sciadv.abe4724

15. Fanning SL, Korngold R, Yang Z, Goldgirsh K, Park S, Zenreich J, et al. Elevated cytokines and chemokines in peripheral blood of patients with SARS-CoV-2 pneumonia treated with high-titer convalescent plasma. PLoS Pathog. 2021;17: e1010025. doi:10.1371/journal.ppat.1010025

16. Tocilizumab in patients admitted to hospital with COVID-19 (RECOVERY): a randomised, controlled, open-label, platform trial. Lancet (London, England). 2021;397: 1637–1645. doi:10.1016/S0140-6736(21)00676-0

17. Salama C, Han J, Yau L, Reiss WG, Kramer B, Neidhart JD, et al. Tocilizumab in Patients Hospitalized with Covid-19 Pneumonia. N Engl J Med. 2021;384: 20–30. doi:10.1056/NEJMoa2030340

18. Beigel JH, Tomashek KM, Dodd LE, Mehta AK, Zingman BS, Kalil AC, et al. Remdesivir for the Treatment of Covid-19 - Final Report. N Engl J Med. 2020;383: 1813–1826. doi:10.1056/NEJMoa2007764

19. Kalil AC, Patterson TF, Mehta AK, Tomashek KM, Wolfe CR, Ghazaryan V, et al. Baricitinib plus Remdesivir for Hospitalized Adults with Covid-19. N Engl J Med. 2021;384: 795–807. doi:10.1056/NEJMoa2031994

20. Marconi VC, Ramanan A V, de Bono S, Kartman CE, Krishnan V, Liao R, et al. Efficacy and safety of baricitinib for the treatment of hospitalised adults with COVID-19 (COV- BARRIER): a randomised, double-blind, parallel-group, placebo-controlled phase 3 trial. Lancet Respir Med. 2021;9: 1407–1418. doi:10.1016/S2213-2600(21)00331-3

21. Hammond J, Leister-Tebbe H, Gardner A, Abreu P, Bao W, Wisemandle W, et al. Oral Nirmatrelvir for High-Risk, Nonhospitalized Adults with Covid-19. N Engl J Med. 2022;386: 1397–1408. doi:10.1056/NEJMoa2118542

22. Alfinito E, Beccaria M, Ciccarese M. Biosensing Cytokine IL-6: A Comparative Analysis of Natural and Synthetic Receptors. Biosensors. 2020;10. doi:10.3390/bios10090106

23. Bryushkova EA, Skatova VD, Mutovina ZY, Zagrebneva AI, Fomina DS, Kruglova TS, et al. Tocilizumab, netakimab, and baricitinib in patients with mild-to-moderate COVID-19: An observational study. PLoS One. 2022;17: e0273340. doi:10.1371/journal.pone.0273340

24. Iovino L, Thur LA, Gnjatic S, Chapuis A, Milano F, Hill JA. Shared inflammatory pathways and therapeutic strategies in COVID-19 and cancer immunotherapy. J Immunother cancer. 2021;9. doi:10.1136/jitc-2021-002392

25. Bonifácio LP, Ramacciotti E, Agati LB, Vilar FC, Silva ACT da, Louzada Júnior P, et al. Efficacy and safety of Ixekizumab vs. low-dose IL-2 vs. Colchicine vs. standard of care in the treatment of patients hospitalized with moderate-to-critical COVID-19: A pilot randomized clinical trial (STRUCK: Survival Trial Using Cytokine Inhibitors). Rev Soc Bras Med Trop. 2023;56: e0565. doi:10.1590/0037-8682-0565-2022

26. Kasiri H, Ghazaiean M, Rouhani N, Naderi-Behdani F, Ghazaeian M, Ghodssi- Ghassemabadi R. The effects of colchicine on hospitalized COVID-19 patients: A randomized, double-blind, placebo-controlled clinical trial. J Investig Med Off Publ Am Fed Clin Res. 2023;71: 124–131. doi:10.1177/10815589221141815

27. Rodriguez C, de Prost N, Fourati S, Lamoureux C, Gricourt G, N’debi M, et al. Viral genomic, metagenomic and human transcriptomic characterization and prediction of the clinical forms of COVID-19. PLoS Pathog. 2021;17: e1009416. doi:10.1371/journal.ppat.1009416

28. Misset B, Piagnerelli M, Hoste E, Dardenne N, Grimaldi D, Michaux I, et al. Convalescent Plasma for Covid-19-Induced ARDS in Mechanically Ventilated Patients. N Engl J Med. 2023;389: 1590–1600. doi:10.1056/NEJMoa2209502

29. Chen P, Nirula A, Heller B, Gottlieb RL, Boscia J, Morris J, et al. SARS-CoV-2 Neutralizing Antibody LY-CoV555 in Outpatients with Covid-19. N Engl J Med. 2021;384: 229–237. doi:10.1056/NEJMoa2029849

30. Guo Y, Huang L, Zhang G, Yao Y, Zhou H, Shen S, et al. A SARS-CoV-2 neutralizing antibody with extensive Spike binding coverage and modified for optimal therapeutic outcomes. Nat Commun. 2021;12: 2623. doi:10.1038/s41467-021-22926-2

31. Sharma A, Ali M. Case Report: Home-based Management of Severe COVID-19 with Low- dose Tofacitinib. Am J Trop Med Hyg. 2021;105: 1472–1475. doi:10.4269/ajtmh.21-0737

32. Batra R, Whalen W, Alvarez-Mulett S, Gomez-Escobar LG, Hoffman KL, Simmons W, et al. Multi-omic comparative analysis of COVID-19 and bacterial sepsis-induced ARDS. PLoS Pathog. 2022;18: e1010819. doi:10.1371/journal.ppat.1010819

33. Varghese E, Samuel SM, Liskova A, Kubatka P, Büsselberg D. Diabetes and coronavirus (SARS-CoV-2): Molecular mechanism of Metformin intervention and the scientific basis of drug repurposing. PLoS Pathog. 2021;17: e1009634. doi:10.1371/journal.ppat.1009634

34. Wing PAC, Prange-Barczynska M, Cross A, Crotta S, Orbegozo Rubio C, Cheng X, et al. Hypoxia inducible factors regulate infectious SARS-CoV-2, epithelial damage and respiratory symptoms in a hamster COVID-19 model. PLoS Pathog. 2022;18: e1010807. doi:10.1371/journal.ppat.1010807

35. Ewart G, Bobardt M, Bentzen BH, Yan Y, Thomson A, Klumpp K, et al. Post-infection treatment with the E protein inhibitor BIT225 reduces disease severity and increases survival of K18-hACE2 transgenic mice infected with a lethal dose of SARS-CoV-2. PLoS Pathog. 2023;19: e1011328. doi:10.1371/journal.ppat.1011328

36. Gamage AM, Tan K Sen, Chan WOY, Liu J, Tan CW, Ong YK, et al. Infection of human Nasal Epithelial Cells with SARS-CoV-2 and a 382-nt deletion isolate lacking ORF8 reveals similar viral kinetics and host transcriptional profiles. PLoS Pathog. 2020;16: e1009130. doi:10.1371/journal.ppat.1009130

37. Çelik N, Çelik O, Laloğlu E, Özkaya A. The CXCL9/10/11-CXCR3 axis as a predictor of COVID-19 progression: a prospective, case-control study. Rev Soc Bras Med Trop. 2023;56: e01282023. doi:10.1590/0037-8682-0128-2023

38. Calvier L, Drelich A, Hsu J, Tseng C-T, Mina Y, Nath A, et al. Circulating Reelin promotes inflammation and modulates disease activity in acute and long COVID-19 cases. Front Immunol. 2023;14: 1185748. doi:10.3389/fimmu.2023.1185748

39. Singh U, Hernandez KM, Aronow BJ, Wurtele ES. African Americans and European Americans exhibit distinct gene expression patterns across tissues and tumors associated with immunologic functions and environmental exposures. Sci Rep. 2021;11: 9905. doi:10.1038/s41598-021-89224-1

40. Dai M, Zhu S, An Z, You B, Li Z, Yao Y, et al. Dissection of key factors correlating with H5N1 avian influenza virus driven inflammatory lung injury of chicken identified by single- cell analysis. PLoS Pathog. 2023;19: e1011685. doi:10.1371/journal.ppat.1011685

41. Files DC, Tacke F, O’Sullivan A, Dorr P, Ferguson WG, Powderly WG. Rationale of using the dual chemokine receptor CCR2/CCR5 inhibitor cenicriviroc for the treatment of COVID-19. PLoS Pathog. 2022;18: e1010547. doi:10.1371/journal.ppat.1010547

42. Gao P, Zhou X-Y, Yashiro-Ohtani Y, Yang Y-F, Sugimoto N, Ono S, et al. The unique target specificity of a nonpeptide chemokine receptor antagonist: selective blockade of two Th1 chemokine receptors CCR5 and CXCR3. J Leukoc Biol. 2003;73: 273–280. doi:10.1189/jlb.0602269

43. Baba M, Nishimura O, Kanzaki N, Okamoto M, Sawada H, Iizawa Y, et al. A small-molecule, nonpeptide CCR5 antagonist with highly potent and selective anti-HIV-1 activity. Proc Natl Acad Sci U S A. 1999;96: 5698–5703. doi:10.1073/pnas.96.10.5698

44. Takama Y, Miyagawa S, Yamamoto A, Firdawes S, Ueno T, Ihara Y, et al. Effects of a calcineurin inhibitor, FK506, and a CCR5/CXCR3 antagonist, TAK-779, in a rat small intestinal transplantation model. Transpl Immunol. 2011;25: 49–55. doi:10.1016/j.trim.2011.04.003

45. Ni J, Zhu Y-N, Zhong X-G, Ding Y, Hou L-F, Tong X-K, et al. The chemokine receptor antagonist, TAK-779, decreased experimental autoimmune encephalomyelitis by reducing inflammatory cell migration into the central nervous system, without affecting T cell function. Br J Pharmacol. 2009;158: 2046–2056. doi:10.1111/j.1476-5381.2009.00528.x

46. Kishi J, Nishioka Y, Kuwahara T, Kakiuchi S, Azuma M, Aono Y, et al. Blockade of Th1 chemokine receptors ameliorates pulmonary granulomatosis in mice. Eur Respir J. 2011;38: 415–424. doi:10.1183/09031936.00070610

47. Tokuyama H, Ueha S, Kurachi M, Matsushima K, Moriyasu F, Blumberg RS, et al. The simultaneous blockade of chemokine receptors CCR2, CCR5 and CXCR3 by a non-peptide chemokine receptor antagonist protects mice from dextran sodium sulfate-mediated colitis. Int Immunol. 2005;17: 1023–1034. doi:10.1093/intimm/dxh284

48. Akashi S, Sho M, Kashizuka H, Hamada K, Ikeda N, Kuzumoto Y, et al. A novel small-molecule compound targeting CCR5 and CXCR3 prevents acute and chronic allograft rejection. Transplantation. 2005;80: 378–384. doi:10.1097/01.tp.0000166338.99933.e1

49. Tan MCB, Goedegebuure PS, Belt BA, Flaherty B, Sankpal N, Gillanders WE, et al. Disruption of CCR5-dependent homing of regulatory T cells inhibits tumor growth in a murine model of pancreatic cancer. J Immunol. 2009;182: 1746–1755. doi:10.4049/jimmunol.182.3.1746

50. Marella M, Chabry J. Neurons and astrocytes respond to prion infection by inducing microglia recruitment. J Neurosci Off J Soc Neurosci. 2004;24: 620–627. doi:10.1523/JNEUROSCI.4303-03.2004

51. Rawle DJ, Le TT, Dumenil T, Yan K, Tang B, Nguyen W, et al. ACE2-lentiviral transduction enables mouse SARS-CoV-2 infection and mapping of receptor interactions. PLoS Pathog. 2021;17: e1009723. doi:10.1371/journal.ppat.1009723

52. Yinda CK, Port JR, Bushmaker T, Offei Owusu I, Purushotham JN, Avanzato VA, et al. K18-hACE2 mice develop respiratory disease resembling severe COVID-19. PLoS Pathog. 2021;17: e1009195. doi:10.1371/journal.ppat.1009195

53. Kumari P, Rothan HA, Natekar JP, Stone S, Pathak H, Strate PG, et al. Neuroinvasion and Encephalitis Following Intranasal Inoculation of SARS-CoV-2 in K18-hACE2 Mice. Viruses. 2021;13. doi:10.3390/v13010132

54. Bishop CR, Dumenil T, Rawle DJ, Le TT, Yan K, Tang B, et al. Mouse models of COVID- 19 recapitulate inflammatory pathways rather than gene expression. PLoS Pathog. 2022;18: e1010867. doi:10.1371/journal.ppat.1010867

55. Chernov AS, Minakov AA, Kazakov VA, Rodionov M V, Rybalkin IN, Vlasik TN, et al. A new mouse unilateral model of diffuse alveolar damage of the lung. Inflamm Res Off J Eur Histamine Res Soc . [et al]. 2022;71: 627–639. doi:10.1007/s00011-022-01568-0

56. Rondovic G, Djordjevic D, Udovicic I, Stanojevic I, Zeba S, Abazovic T, et al. From Cytokine Storm to Cytokine Breeze: Did Lessons Learned from Immunopathogenesis Improve Immunomodulatory Treatment of Moderate-to-Severe COVID-19? Biomedicines. 2022;10. doi:10.3390/biomedicines10102620

57. Bi Z, Hong W, Que H, He C, Ren W, Yang J, et al. Inactivated SARS-CoV-2 induces acute respiratory distress syndrome in human ACE2-transgenic mice. Signal Transduct Target Ther. 2021;6: 439. doi:10.1038/s41392-021-00851-6

58. Andreakos E, Papadaki M, Serhan CN. Dexamethasone, pro-resolving lipid mediators and resolution of inflammation in COVID-19. Allergy. Denmark; 2021. pp. 626–628. doi:10.1111/all.14595

59. Ahmed MH, Hassan A. Dexamethasone for the Treatment of Coronavirus Disease (COVID- 19): a Review. SN Compr Clin Med. 2020;2: 2637–2646. doi:10.1007/s42399-020-00610-8

60. Horby P, Lim WS, Emberson JR, Mafham M, Bell JL, Linsell L, et al. Dexamethasone in Hospitalized Patients with Covid-19. N Engl J Med. 2021;384: 693–704. doi:10.1056/NEJMoa2021436

61. Pinzón MA, Ortiz S, Holguín H, Betancur JF, Cardona Arango D, Laniado H, et al. Dexamethasone vs methylprednisolone high dose for Covid-19 pneumonia. PLoS One. 2021;16: e0252057. doi:10.1371/journal.pone.0252057

62. Bouadma L, Mekontso-Dessap A, Burdet C, Merdji H, Poissy J, Dupuis C, et al. High-Dose Dexamethasone and Oxygen Support Strategies in Intensive Care Unit Patients With Severe COVID-19 Acute Hypoxemic Respiratory Failure: The COVIDICUS Randomized Clinical Trial. JAMA Intern Med. 2022;182: 906–916. doi:10.1001/jamainternmed.2022.2168

63. Wu H, Daouk S, Kebbe J, Chaudry F, Harper J, Brown B. Low-dose versus high-dose dexamethasone for hospitalized patients with COVID-19 pneumonia: A randomized clinical trial. PLoS One. 2022;17: e0275217. doi:10.1371/journal.pone.0275217

64. Wyler E, Adler JM, Eschke K, Teixeira Alves G, Peidli S, Pott F, et al. Key benefits of dexamethasone and antibody treatment in COVID-19 hamster models revealed by single- cell transcriptomics. Mol Ther. 2022;30: 1952–1965. doi:10.1016/j.ymthe.2022.03.014

65. Meng Q-F, Tai W, Tian M, Zhuang X, Pan Y, Lai J, et al. Inhalation delivery of dexamethasone with iSEND nanoparticles attenuates the COVID-19 cytokine storm in mice and nonhuman primates. Sci Adv. 2023;9: eadg3277. doi:10.1126/sciadv.adg3277

66. Yuan L, Zhou M, Ma J, Liu X, Chen P, Zhu H, et al. Dexamethasone ameliorates severe pneumonia but slightly enhances viral replication in the lungs of SARS-CoV-2-infected Syrian hamsters. Cellular & molecular immunology. China; 2022. pp. 290–292. doi:10.1038/s41423-021-00793-7

67. Xu T, Qiao J, Zhao L, He G, Li K, Wang J, et al. Effect of dexamethasone on acute respiratory distress syndrome induced by the H5N1 virus in mice. Eur Respir J. 2009;33: 852–860. doi:10.1183/09031936.00130507

68. Xu X, Han M, Li T, Sun W, Wang D, Fu B, et al. Effective treatment of severe COVID-19 patients with tocilizumab. Proc Natl Acad Sci U S A. 2020;117: 10970–10975. doi:10.1073/pnas.2005615117

69. Meoni G, Ghini V, Maggi L, Vignoli A, Mazzoni A, Salvati L, et al. Metabolomic/lipidomic profiling of COVID-19 and individual response to tocilizumab. PLoS Pathog. 2021;17: e1009243. doi:10.1371/journal.ppat.1009243

70. Rosas IO, Bräu N, Waters M, Go RC, Hunter BD, Bhagani S, et al. Tocilizumab in Hospitalized Patients with Severe Covid-19 Pneumonia. N Engl J Med. 2021;384: 1503– 1516. doi:10.1056/NEJMoa2028700

71. Stone JH, Frigault MJ, Serling-Boyd NJ, Fernandes AD, Harvey L, Foulkes AS, et al. Efficacy of Tocilizumab in Patients Hospitalized with Covid-19. N Engl J Med. 2020;383: 2333–2344. doi:10.1056/NEJMoa2028836

72. Mehta M, Purpura LJ, McConville TH, Neidell MJ, Anderson MR, Bernstein EJ, et al. What about tocilizumab? A retrospective study from a NYC Hospital during the COVID-19 outbreak. PLoS One. 2021;16: e0249349. doi:10.1371/journal.pone.0249349

73. Veiga VC, Prats JAGG, Farias DLC, Rosa RG, Dourado LK, Zampieri FG, et al. Effect of tocilizumab on clinical outcomes at 15 days in patients with severe or critical coronavirus disease 2019: randomised controlled trial. BMJ. 2021;372: n84. doi:10.1136/bmj.n84

74. Gu T, Zhao S, Jin G, Song M, Zhi Y, Zhao R, et al. Cytokine Signature Induced by SARS- CoV-2 Spike Protein in a Mouse Model. Front Immunol. 2020;11: 621441. doi:10.3389/fimmu.2020.621441

75. Hu J, Feng X, Valdearcos M, Lutrin D, Uchida Y, Koliwad SK, et al. Interleukin-6 is both necessary and sufficient to produce perioperative neurocognitive disorder in mice. Br J Anaesth. 2018;120: 537–545. doi:10.1016/j.bja.2017.11.096

76. Kamiya N, Kuroyanagi G, Aruwajoye O, Kim HKW. IL6 receptor blockade preserves articular cartilage and increases bone volume following ischemic osteonecrosis in immature mice. Osteoarthr Cartil. 2019;27: 326–335. doi:10.1016/j.joca.2018.10.010

77. Orabona C, Mondanelli G, Pallotta MT, Carvalho A, Albini E, Fallarino F, et al. Deficiency of immunoregulatory indoleamine 2,3-dioxygenase 1in juvenile diabetes. JCI insight. 2018;3. doi:10.1172/jci.insight.96244

78. Wu R, Liu X, Yin J, Wu H, Cai X, Wang N, et al. IL-6 receptor blockade ameliorates diabetic nephropathy via inhibiting inflammasome in mice. Metabolism. 2018;83: 18–24. doi:10.1016/j.metabol.2018.01.002

79. Lokau J, Kleinegger F, Garbers Y, Waetzig GH, Grötzinger J, Rose-John S, et al. Tocilizumab does not block interleukin-6 (IL-6) signaling in murine cells. PLoS One. 2020;15: e0232612. doi:10.1371/journal.pone.0232612

80. Patterson BK, Seethamraju H, Dhody K, Corley MJ, Kazempour K, Lalezari J, et al. CCR5 inhibition in critical COVID-19 patients decreases inflammatory cytokines, increases CD8 T-cells, and decreases SARS-CoV2 RNA in plasma by day 14. Int J Infect Dis IJID Off Publ Int Soc Infect Dis. 2021;103: 25–32. doi:10.1016/j.ijid.2020.10.101

81. Chang XL, Wu HL, Webb GM, Tiwary M, Hughes C, Reed JS, et al. CCR5 Receptor Occupancy Analysis Reveals Increased Peripheral Blood CCR5+CD4+ T Cells Following Treatment With the Anti-CCR5 Antibody Leronlimab. Front Immunol. 2021;12. doi:10.3389/fimmu.2021.794638

82. Agresti N, Lalezari JP, Amodeo PP, Mody K, Mosher SF, Seethamraju H, et al. Disruption of CCR5 signaling to treat COVID-19-associated cytokine storm: Case series of four critically ill patients treated with leronlimab. J Transl Autoimmun. 2021;4: 100083. doi:10.1016/j.jtauto.2021.100083

83. Gaylis NB, Ritter A, Kelly SA, Pourhassan NZ, Tiwary M, Sacha JB, et al. Reduced Cell Surface Levels of C-C Chemokine Receptor 5 and Immunosuppression in Long Coronavirus Disease 2019 Syndrome. Clin Infect Dis an Off Publ Infect Dis Soc Am. 2022;75: 1232– 1234. doi:10.1093/cid/ciac226

84. Okamoto M, Toyama M, Baba M. The chemokine receptor antagonist cenicriviroc inhibits the replication of SARS-CoV-2 in vitro. Antiviral Res. 2020;182: 104902. doi:10.1016/j.antiviral.2020.104902

85. Thompson M, Saag M, DeJesus E, Gathe J, Lalezari J, Landay AL, et al. A 48-week randomized phase 2b study evaluating cenicriviroc versus efavirenz in treatment-naive HIV- infected adults with C-C chemokine receptor type 5-tropic virus. AIDS. 2016;30: 869–878. doi:10.1097/QAD.0000000000000988

86. Friedman SL, Ratziu V, Harrison SA, Abdelmalek MF, Aithal GP, Caballeria J, et al. A randomized, placebo-controlled trial of cenicriviroc for treatment of nonalcoholic steatohepatitis with fibrosis. Hepatology. 2018;67: 1754–1767. doi:10.1002/hep.29477

87. Ratziu V, Sanyal A, Harrison SA, Wong VW-S, Francque S, Goodman Z, et al. Cenicriviroc Treatment for Adults With Nonalcoholic Steatohepatitis and Fibrosis: Final Analysis of the Phase 2b CENTAUR Study. Hepatology. 2020;72: 892–905. doi:10.1002/hep.31108

88. O’Halloran JA, Ko ER, Anstrom KJ, Kedar E, McCarthy MW, Panettieri RAJ, et al. Abatacept, Cenicriviroc, or Infliximab for Treatment of Adults Hospitalized With COVID- 19 Pneumonia: A Randomized Clinical Trial. JAMA. 2023;330: 328–339. doi:10.1001/jama.2023.11043

89. Shamsi A, Mohammad T, Anwar S, AlAjmi MF, Hussain A, Rehman MT, et al. Glecaprevir and Maraviroc are high-affinity inhibitors of SARS-CoV-2 main protease: possible implication in COVID-19 therapy. Biosci Rep. 2020;40. doi:10.1042/BSR20201256

90. Patterson BK, Yogendra R, Guevara-Coto J, Mora-Rodriguez RA, Osgood E, Bream J, et al. Case series: Maraviroc and pravastatin as a therapeutic option to treat long COVID/Post- acute sequelae of COVID (PASC). Front Med. 2023;10: 1122529. doi:10.3389/fmed.2023.1122529

91. Risner KH, Tieu K V, Wang Y, Getz M, Bakovic A, Bhalla N, et al. Maraviroc inhibits SARS-CoV-2 multiplication and s-protein mediated cell fusion in cell culture. bioRxiv : the preprint server for biology. United States; 2022. doi:10.1101/2020.08.12.246389

92. Sharapova TN, Romanova EA, Chernov AS, Minakov AN, Kazakov VA, Kudriaeva AA, et al. Protein PGLYRP1/Tag7 Peptides Decrease the Proinflammatory Response in Human Blood Cells and Mouse Model of Diffuse Alveolar Damage of Lung through Blockage of the TREM-1 and TNFR1 Receptors. Int J Mol Sci. 2021;22. doi:10.3390/ijms222011213

